# Annotation-agnostic discovery of associations between novel gene isoforms and phenotypes

**DOI:** 10.1101/2022.12.02.518787

**Authors:** Kristján Eldjárn Hjörleifsson, Lior Pachter, Páll Melsted

## Abstract

We present a novel method for associating phenotypes with RNA expression, that can identify expression associations resulting from a wide variety of underlying transcriptional and post-transcriptional events, without relying on annotations of these events. We show that we can reliably detect, *de novo*, phenotypically relevant transcriptional structures

## Introduction

The quantification of RNA reads is a key step in most analyses of RNA-seq data (Kukurba and Montgomery, 2015). Current quantification methods rely on annotations of the organisms’ transcriptomes, which may be incomplete or nonexistent (Zhang et al. 2020). This may result in data being discarded and can lead to erroneous quantifications. In downstream applications such as eQTL analysis, such errors can propagate and result in missed, or erroneous associations (Saha and Battle, 2018). We present a novel method for associating phenotypes with RNA expression, that can identify expression associations resulting from a wide variety of underlying transcriptional and post-transcriptional events, without requiring a prior annotation of the transcriptome. By constructing a *de Bruijn* graph (Bruijn, de, 1946) of all the reads overlapping a single gene, and pruning away nodes that are likely to be due to sequencing errors, we obtain a representation of the expression of the gene in our cohort. Each expressed isoform constitutes one path through the graph. We then run associations on the expression of each individual node and a phenotype. Should an isoform of the gene associate with the phenotype, there will be a set of nodes in the graph that uniquely identify the isoform, the expression of which also associates with expression of the phenotype. This method enables discovery of novel alternative polyadenylation, exon-skipping, duplications, insertions, deletions, and circular RNA, among other transcriptional and post-transcriptional variations, without prior knowledge of these events. We show that we can reliably reproduce known associations, and detect, *de novo*, phenotypically relevant transcriptional structures.

## Methods

RNA-Seq reads overlapping a genomic region of interest, e.g. a gene, are obtained from a cohort of people for which a phenotype of interest is available. A bi-directed *de Bruijn* Graph (dBG) is constructed, using Bifrost (Holley and Melsted, 2020), from these reads with *k*-mer size *k* = 31 and then compacted such that consecutive *k*-mers with out-degree 1 and in-degree 1 respectively are folded into a single, maximal *unitig*, which is a high-confidence contig. Each path between two unitigs represents distinct ways the corresponding part of the gene might be expressed in the cohort. Each individual’s expression of each unitig in the graph is then quantified.

### Quality filter

The size of the dBG is dependent on several factors: genetic variation, errors introduced in sequencing, the size and relatedness of the cohort, the length and number of different isoforms of the gene, the expression levels of that gene in the tissue of interest, etc. In order to reduce the scope of the problem and thereby reduce the number of association tests performed, we may prune from the graph nodes that are likely present due to noise; either genetic or technical. To that end, all kmers that occur in common transposable elements, such as Arthrobacter luteus (Alu) regions are removed from the graph. Furthermore, the median abundance of isoforms of the gene that are present in an annotation of the target transcriptome are found for each individual in the cohort, and unitigs that are expressed less than 0.5% of the transcriptomic median abundance are deemed to be erroneous. Unitigs with less than 50 counts associated with them are taken to be erroneous if there is another unitig within Hamming distance 1, with more than four times their expression. Finally, tips, i.e. short chains of nodes that are disconnected on one end (Zerbino and Birney, 2008) are removed.

### Associations

Unitig abundances are normalized to sum up to 1 for each individual, in order to capture associations between genotypes and phenotypes on one hand, and isoform-specific expression rather than individual coverage, on the other. The normalized abundances for each unitig respectively, are associated with a qualitative or a qualitative phenotype. Crucially, the expression of sequences from any isoforms containing structural variants, which are not part of any transcriptome annotation, are implicitly associated with the phenotype. For any such novel isoform, there will be a set of subsequences, the expression of which uniquely distinguishes the isoform from other transcripts. Since the relative expression levels of different isoforms of the same gene are not generally independent, the resulting *p*-values for the individual unitigs are aggregated using the Harmonic Mean P-value (HMP) (Wilson, 2019) with weights equal to the log-transformed mean counts normalized to 1, i.e. given a dBG with *N* unitigs with mean counts *u*_*1*_ , ..., *u*_*N*_, the weight for the p value of unitig *i* is 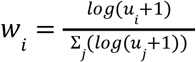 (Yi et al. 2018). Distinct genes are assumed to be expressed independently, and the aggregated p-value is Bonferroni corrected for the number of protein-coding genes in the target transcriptome.

### Simulated differential transcript usage experiments

In order to assess the sensitivity of the method, we simulated bulk RNA-Seq datasets with novel structures and simulated phenotypes that correlated with those novel structures. While methods to generate signals that do not require theoretical models exist (Gerard, 2020), these generate their signal using reads from real RNA-seq datasets. In our case we need to simulate novel isoforms as well the signals correlated with them. Basing the signal simulation on real-world RNA-seq datasets is not tenable in our case, since they would not contain the novel isoform. A protein-coding gene was arbitrarily selected from version 108 of the Ensembl annotation of the GRCh38 assembly (Cunningham et al., 2022). An alternative transcriptome *T*_*alt*_ was generated from the reference transcriptome *T*_*ref*_ by adding a novel isoform based on an existing isoform, but containing a duplication, exon skipping, alternative polyadenylation, or circular structure not present in the original. A cohort of size 1000 was created, 10 of which were chosen to express the novel structure, for a prevalence of 1%. Single-ended reads overlapping the chosen protein-coding gene were generated for all 1000 individuals in the cohort using BBMap (Bushnell, 2014). For the affected cohort, the reads were generated from *T*_*alt*_ whereas for the remaining 990 wild-type individuals the reads were generated from *T*_*ref*_. Various different rates for SNPs and indels were used in order to assess robustness to noise. Qualitative phenotypes were obtained by assigning the affected individuals expressing the novel structure the phenotype 1 and others 0.

Varying the allele frequency of SNPs and indels gives us an idea of the level of robustness to biological noise. A number of simulations were run, with the numbers of reads per individual varying from 1 to 100, in order to assess the method’s sensitivity to coverage, with 25 reads per gene taken to be a reasonable coverage per Svensson et al. (2017), and assuming 19,116 protein-coding genes in the transcriptome (Piovesan et al., 2019). Simulating different levels of coverage shows us the sensitivity of the method as a function of the expected number of reads overlapping an identifying sequence of the novel isoform, given by

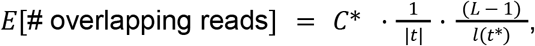

Where *C\** denotes the total number of reads in affected individuals, *t* denotes the set of isoforms, *L* denotes the expected read length, and *l*(*t\**) denotes the length of the novel isoform. Expected coverage of an identifying sequence in the range of [1, 50] was simulated and quantified using the method [Figure 2.A]. Technical noise was simulated by adding or subtracting from each unitig for each individual a number of reads drawn from a Poisson distribution with parameter λ = µ · ξ, where µ is the mean expression over all individuals and unitigs, and ξ is a scaling factor. Values of ξ ∈ [0. 0*1*, 0. *1*] were simulated [Figure 2.C] to assess the robustness of the method w.r.t. technical noise. Genetic noise was simulated by varying the Single Nucleotide Polymorphism (SNP) rate *r*_*SNP*_ [Supplementary appendix]. Values of *r*_*SNP*_ ∈ [0. 0*5*, 0. *1*, 0. *5*] were simulated [Figure 2.B] to assess the robustness of the method w.r.t. genetic noise. For each set of parameters, 1000 simulated experiments were run and the proportion of experiments where a signal was discovered was reported. Associations were performed using a Wilcoxon rank-sum test, and Bonferroni-corrected for 19,116 protein-coding genes (Piovesan et al., 2019), yielding a significance threshold of p < 2.5e-6.

**Figure 1.**
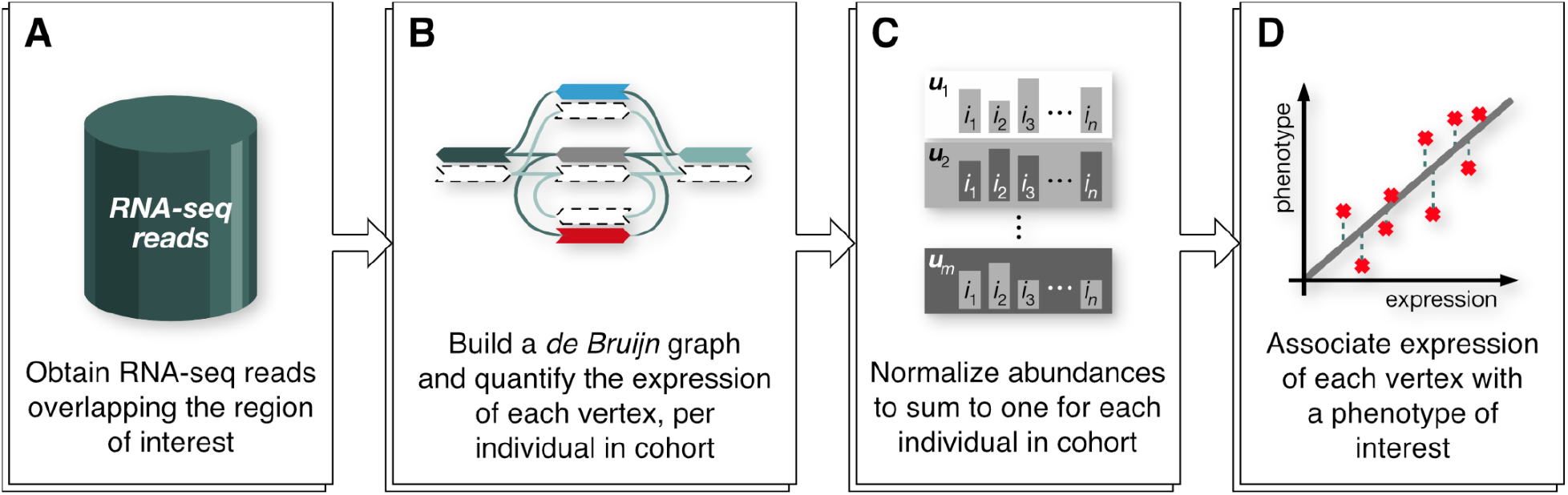
The annotation-agnostic association process consists of four steps. **A**. We obtain a dataset of RNA-seq reads from a cohort of individuals, overlapping the genomic region of interest. **B**. A *de Bruijn* graph is constructed from the dataset, and the individuals’ expression of each vertex is quantified. An optional pruning step can remove vertices that are likely to be erroneous (either due to genetic or technical noise) in order to reduce the number of association targets [methods]. **C**. The vertex abundances are normalized such that the individuals’ expression sums to one, respectively. **D**. The normalized expression of each vertex is associated with a phenotype of interest, and the *p*-values of the associations are aggregated using the harmonic mean *p*-value [Methods].

**Figure 2.**
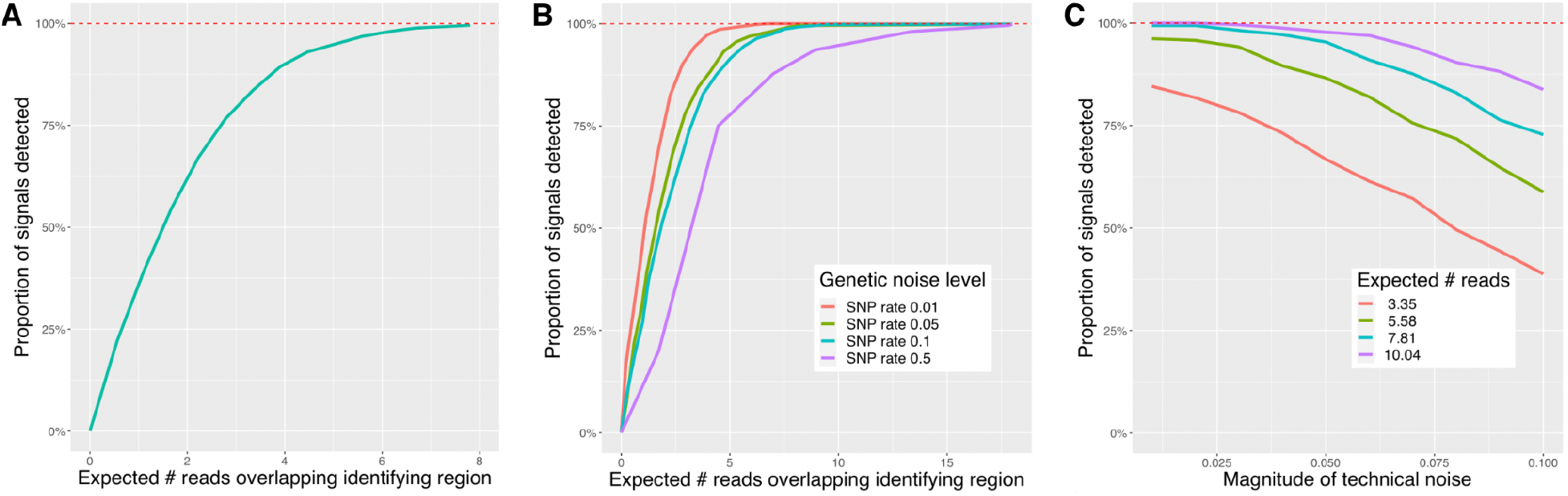
The proportion of 1000 simulated annotation-agnostic association experiments in which a ground truth association between expression of a novel isoform, containing a duplication, and a phenotype, was detected. **A**. To detect a signal in an experiment with no genetic or technical noise, it was sufficient for an expected 5 reads to overlap an identifying region of the gene. **B**. Even with a SNP rate of 0.5 [Methods], we detected over 95% of all signals, with an expected 11 reads overlapping an identifying region of the gene. Note however, that even with quality filters in place [Methods] the noise adds extraneous vertices to the graph, the expression of which must also be associated with the phenotype, resulting in a large number of computations. **C**. With an expected 10 reads overlapping an identifying region of the gene, we can reliably detect signals from simulations with technical noise [Methods] of magnitude up to 7% of the mean expression in the cohort.

The simulated reads were quantified using kallisto (Bray et al. 2016), using an index constructed from *T*_*ref*_, in order to attempt to discover associations between those abundances and the target phenotype.

## Code availability

The method was implemented in C++ using the Bifrost library (Holley and Melsted, 2020) for the construction and maintenance of de Bruijn Graphs. The source code is available under GPLv3 and can be downloaded from https://github.com/pachterlab/AAQuant. The simulation framework is available at https://github.com/pachterlab/HPM_2022.

## Results

### Sensitivity analysis

As evident in Figure 2.A, we can reliably recover associations between the expression of a novel isoform and a qualitative phenotype, even with low expected coverage of the identifying locus. These associations are not discovered when using transcript abundances from kallisto, using version 104 of the Ensembl annotation of GRCh38. Furthermore, Figure 2.B shows that even for high levels of genetic noise, we can still detect these associations, due to the pruning of low quality vertices from the graph. Lastly, per Figure 2.C, we can reliably recover the associations with high levels of instrumental noise, and the robustness to noise is relative to the expected number of reads overlapping a distinguishing region. Combining these three measures of robustness, we are able to detect associations 97.5% of all associations between the expression of a novel isoform and a qualitative phenotype, using reasonable parameters for a real-world experiment, e.g. 25 reads per individual, which yields an expected 13.9 reads overlapping a distinguishing region, SNP rate of 1%, and noise with a magnitude of 5% of the mean unitig expression.

## Discussion

AbundanceDBG enables annotation-agnostic discovery of associations between relative abundances of kmers in gene transcripts on one hand, and qualitative and quantitative phenotypes on the other. It does so in a memory and computationally efficient way by processing unambiguous, overlapping *k*-mers together, and by leveraging minimizers for lookup in the underlying graph. The ability to detect transcriptional and post-transcriptional events, without prior knowledge of those events is useful for discovering expression associations in instances where transcriptome annotations are incomplete or nonexistent. Firstly, we have shown that we can discover association between novel isoforms and a phenotype, without prior knowledge of the isoforms. Secondly, using simulated experiments, we have shown that we can reliably detect associations not found by state-of-the-art RNA-seq quantification methods. We have furthermore demonstrated that our discovery of the associations is robust to genetic and instrumental noise. However, the method does not attribute meaning to the associated sequence. Having discovered an association between a phenotype and a sequence, it must then be aligned against the genome to identify the transcriptional or post-transcriptional events that yielded the sequence. As an illustration, if the associated sequence was produced by a duplication in the gene, it remains to be determined where in the sequence the duplication splice junction is, and the loci of the sequences on either side of the junction.

## Supplementary appendix

### Read simulations

Single-ended reads of length 150 basepairs of the gene COX5B were simulated using BBMap (Bushnell, 2014). The reads were simulated from two different transcriptomes; one containing a novel duplication of length 55, and the other only containing the isoforms present in the GRCh38 genome annotation. SNPs were simulated such that for a rate of *r*_*SNP*_, each read has a (100 × *r*_*snp*_)% chance of containing a single SNP, (100 × *r*_*SNP*_)^2^ % chance of containing two SNPs, &c. Reads for SNP rates *r*_*SNP*_ ∈ [0. 05 , 0. 1 , 0. 5 , 1 , 1 . 5 , 2, 2. 5] were simulated.

